# A NOVEL METHOD FOR BLOOD DETECTION USING FLUORESCENT DYE

**DOI:** 10.1101/2022.12.04.519063

**Authors:** Kamayani Vajpayee, Preet Desai, Hirak Ranjan Dash, Ritesh Kumar Shukla

## Abstract

Detection of the body fluid present at a crime scene is essential for any forensic investigation. Amongst all the body fluids (sweat, semen, vaginal fluid, saliva, etc.,) blood is the most frequently encountered evidence at the crime scene. Currently, reagents like benzidine, ortho-toluidine, tetramethylbenzidine, phenolphthalein, leucomalachite green, luminol, and fluorescein are used to screen the presence of blood on different surfaces (porous/nonporous). Majority of these tests are based on colorimetric changes owing to the nature of hemoglobin to catalyze the oxidation of chromogenic compounds. Apart from aiding the investigation, these reagents show toxic behavior (DNA damage, carcinogenic, etc.) and false-positive results. Hence, to circumvent this issue, the present study attempts to develop a state-of-art methodology for preliminary blood detection and screening using fluorescein-derived 2’,7’-dichlorofluorescein di-acetate (DCFDA) dye. It is hereby proposed that the fluorescein-based dye can successfully detect blood and bloodstains aged up to 20 days. Moreover, supplemental experiments have suggested that the dye doesn’t interfere with DNA integrity-causing any damage to DNA. Parallelly, no false-positive reactions have been observed as tested against similar-looking substances.

## Introduction

Identification is pivotal to the criminal justice system. Shreds of evidence found at the crime scene play a decisive role in identification [1]. These evidences are tell-tales of the crime scene. They speak of how the crime happened and who all were involved in its commencement. It links the scene of crime (SoC) with the victim(s) and culprit(s), further aiding the crime scene reconstruction [2]. The evidences found at the crime scene can be of varied types. They are broadly categorized as *Physical* and *Biological evidences*-based on their origin. Both of these are ad rem to any investigation [3]. However, biological evidence has gotten the scientific community’s profound attention in the past decade. This can be attributed to their properties which are regularly exploited for individualization purposes in forensic casework like DNA profiling technology [4]. Forensic professionals have been attempting to improve identification and individualization approaches using the enormous power of biological evidence. These attempts thus paved the way for the emergence of several branches in the field of forensic science, namely, forensic anthropology, fingerprint analysis, forensic serology, etc.

Forensic serology can be defined as the branch of forensic science which uses immunological and biochemical procedures to determine the presence of a body fluid such as blood, saliva, semen, urine, and other fluids or tissue sample acquired during a criminal investigation primarily for identification purposes [5,6].

Determining whether a bodily fluid is present and then identifying it permits the sample to undergo additional laboratory testing, such as DNA analysis, which is a critical stage in various investigations. However, this is not always an easy task because many body fluid stains are either seem similar to other fluids/substances or invisible to the human eye. Even if a forensic investigator deems the identity of a stain is evident, definitive proof is required before the evidence may be utilized in court to show or refute a fact in a case. This becomes most important when dealing with the possibility of combinations. Multiple body fluids from two or more donors could be present in a stain. Physical examinations on these questioned stains allow the crime scene/forensic experts to identify a fluid or establish the absence of one, both of which are important in a case. Each of these fluids has one or more presumptive screening tests, some of which have confirmatory tests to prove their presence and identify the species they originated from. [4].

Blood is the most commonly encountered biological evidence [7]. Blood and bloodstains can be found in forensic casework like homicide, assault, mass disasters, etc. [8]. Human blood mainly consists of erythrocytes (red blood corpuscles), leukocytes (white blood corpuscles), and platelets suspended in a protein-rich fluid known as blood plasma. Red blood cells (RBCs) make up 40–45 percent of total blood volume; leukocytes and platelets each make up less than 1% of total blood volume. The only cells in the blood that contain DNA are leukocytes [9]. RBCs consist of an iron-containing metalloprotein molecule called hemoglobin (Hb) that is structurally arranged where the heme (iron) molecule at the center is surrounded by four polypeptide chains (two alpha chains and two beta chains) [10].

Several preliminary and confirmatory tests have been designed and validated to identify blood and bloodstains. The tests depend on the catalytic oxidation-reduction reaction between chemical and heme molecule (present inside the blood) in the presence of hydrogen peroxide (H_2_O_2_), which are typically presumptive and have little specificity or sensitivity. In contrast, confirmatory tests rely upon microscopic or immunological examinations [11].

Forensic experts currently employ these assays and tests; nevertheless, several limitations are associated. These assays are nonspecific and tend to show false positive results with other substances like chemical oxidants and fruit/vegetable peroxidases [12,13]. In addition, some of these tests, as given in Table 1, have been reported to be carcinogenic [13,14], thus, posing an occupational exposure to the experts dealing with them regularly. They either use UV light to expose the stain or use reagents generating reactive oxygen species (ROS), which conflicts with the downstream assay like DNA profiling [15,16]. Due to their sensitivity, certain assays, including luminol, have also been shown to consume samples that were already scarce completely [16-19]. Advancements in science and technology have introduced spectroscopic methods, such as UV–vis absorption for bloodstain screening and identification [20]. Although the techniques are sensitive, conditions like water immersion, sunshine exposure, heating, and rust can affect spectral readings [10,12]. Other limitations include lengthy development time, additional pre-saturation steps, the requirement of skilled labor, and, most importantly, cost [12].

**Table 1.** Currently available blood screening tests with their limitations.

| S.No. | Blood Screening Tests | Limitations | References |
| --- | --- | --- | --- |
| 1. | Alternating Light Source | Damages DNA | [18] |
| 2. | Grodsky formulation | Damages DNA | [15] |
| 3. | Benzidine test (Alders' Test) | False-positive, Carcinogenic | [12,13] |
| 4. | Kastle-Meyer Test (Phenolphthalein Test) | False Positive | [12] |
| 5. | Leucomalachite green (LMG) Test | Specificity | [20] |

Thus, the abovementioned limitations opened the doors to developing novel blood presumptive tests. Given this, 2’,7’-dichlorofluorescein di-acetate (DCFDA), a fluorescein derivative non-fluorescent dye, has shown to be an alternative to all the currently used reagents.

DCFDA dye, on the one hand, detects blood and bloodstains with utmost accuracy and sensitivity. On the other hand, it does not cause potential damage to the genetic material, i.e., DNA. Therefore, a cutting-edge technique for preliminary blood detection and screening employing the dye 2’,7’-dichlorofluorescein di-acetate (DCFDA) derived from fluorescein has been devised to overcome this challenge.

## Materials and Methods

### Samples and their preparation, quality control and Ethical statement

Peripheral liquid blood was collected from the volunteers after obtaining their written informed consent. The study has been conducted with the approval of the Ethics committee of B.J. Medical College & Civil Hospital, Ahmedabad, Gujarat, India (Ref. No. — EC/Approval/109/07.04.2017). The blood samples collected in K_2_EDTA vials were kept at 4_ till further use.

Commercially available 2, 7-dichloro-fluorescein-di-acetate was purchased from Sigma Aldrich, USA. Phosphate Buffer Saline (PBS) powder was purchased from Sigma Aldrich, USA. Glass slides with frosted end and coverslips were purchased from Blue Label, India. Dimethyl sulfoxide (DMSO) was purchased from Merck Limited, India.

Pomegranate and beetroot were procured from the local market, and the fresh juice was extracted from them. The juices were kept separately in vials and were stored at 4° C until use. Tomato Ketchup (Kissan fresh, Hindustan Unilever Limited, India) was purchased from the local shop and was refrigerated until use.

Fluorescent microscope (Leica DM 2500) was purchased from Leica Microsystems, Germany.

### Preparation of Phosphate Buffer Saline

Phosphate buffer saline solution (Ca ^2+^, Mg ^2+^ free) was prepared using Phosphate Buffer Saline (PBS) powder (Sigma Aldrich, USA.). 9.6 grams of the powdered reagent was added to 990 milliliters of ultra-purified milli-Q water (Merck, Millipore systems). A pH of 7.4 was maintained using 1 N Hydrochloric acid (HCl) (Sigma Aldrich, USA). The final volume was adjusted to 1000 milliliters using milli-Q water.

#### Preparation of DCFDA solution

Stock and working solutions of DCFDA dye were prepared. The stock solution (10mM) of DCF-DA was prepared by adding 5 mg of DCF-DA powder to 1 ml of DMSO. Further, the working solution of 20μM was prepared by adding 500μl of the stock solution into 4.5 ml of PBS.

#### Reaction with Fresh, pureblood sample

To carryout the reactions in the entire study, Glass slides were used. The slides were of two types: transparent surface (glass slides) and dark/black bottom surface. 15μl of pure blood sample was taken on both surfaces. It was then allowed to dry at normal room temperature. To this, 25 μl of working solution (20μM) was added. The colorimetric change was instantly observed by unaided eyes,The fluorimetric analysis was then done under a fluorescent microscope (Leica DM 2500) at 490/535 nm (N2.1 filter).

#### Reaction with Fresh diluted blood

Blood dilutions of 10x, 100x, 1000x, 10,000x and 1,00,000x were prepared in PBS. 15μl of each diluted sample was taken on the surface of the glass slide (transparent and black bottom surface). It was further allowed to dry at normal room temperature. After that, 25 μl of the working solution (20μM) was added. The reaction mixture was then analyzed under a fluorescent microscope.

#### Reaction with Aged Blood samples

Pure and diluted blood samples were kept from 24 hours to 30 days. Day-wise analysis of the samples with DCFDA solution was performed where 15μl of samples (kept for different periods) were taken on separate, clean, and sterilized surfaces. 25 μl of the working solution was added, followed by observations under the fluorescent microscope.

#### Reaction with similar-looking stains

Previous studies have reported that the reagents used for preliminary blood screening tend to give false-positive tests. Thus, to further validate the proposed method against false-positive results, similar-looking stains - Beetroot juice, Tomato Ketchup, and Pomegranate juice were tested for their reactivity with DCFDA solution on the same surfaces.

Each specimen was subjected to above mentioned DCFDA protocol. The observations were made for fresh and aged samples for up to 30 days.

#### Fluorescent Spectroscopy

Fluorescence spectra were acquired using a Leica DM 2500 (Leica Microsystems, Germany) with a 56°C excitation incidence angle, 5 nm excitation and emission slits. At 485 nm and 535 nm, the excitation/emission spectra of DCFDA were detected. The samples were evaluated in duplicates.

## Results

To circumvent the harmful effects of peroxide and carcinogenic dyes used in blood presumptive tests, 2,7 dichloro-fluorescein-di-acetate (DCF-DA) has been tested against bloodstain and similar-looking stains like tomato ketchup, beetroot juice, and pomegranate juice for its efficiency in blood screening.

### The reaction of DCF-DA with Blood

As discussed above, fresh and aged blood samples were treated with the proposed non-carcinogenic fluorescent dye-DCFDA for up to 30 days. It was observed that the reaction between the DCFDA and blood in the presence of blood esterases results in the formation of fluorescent spectra. The fluorescence spectra so formed (Bright green halo) were observed at 485 nm / 535 nm.

Similarly, the fluorescent spectra were observed for the samples aged up to 20 days (Figure 1). However, samples kept for more than 20 days failed to produce fluorescence.

**Figure 1:**
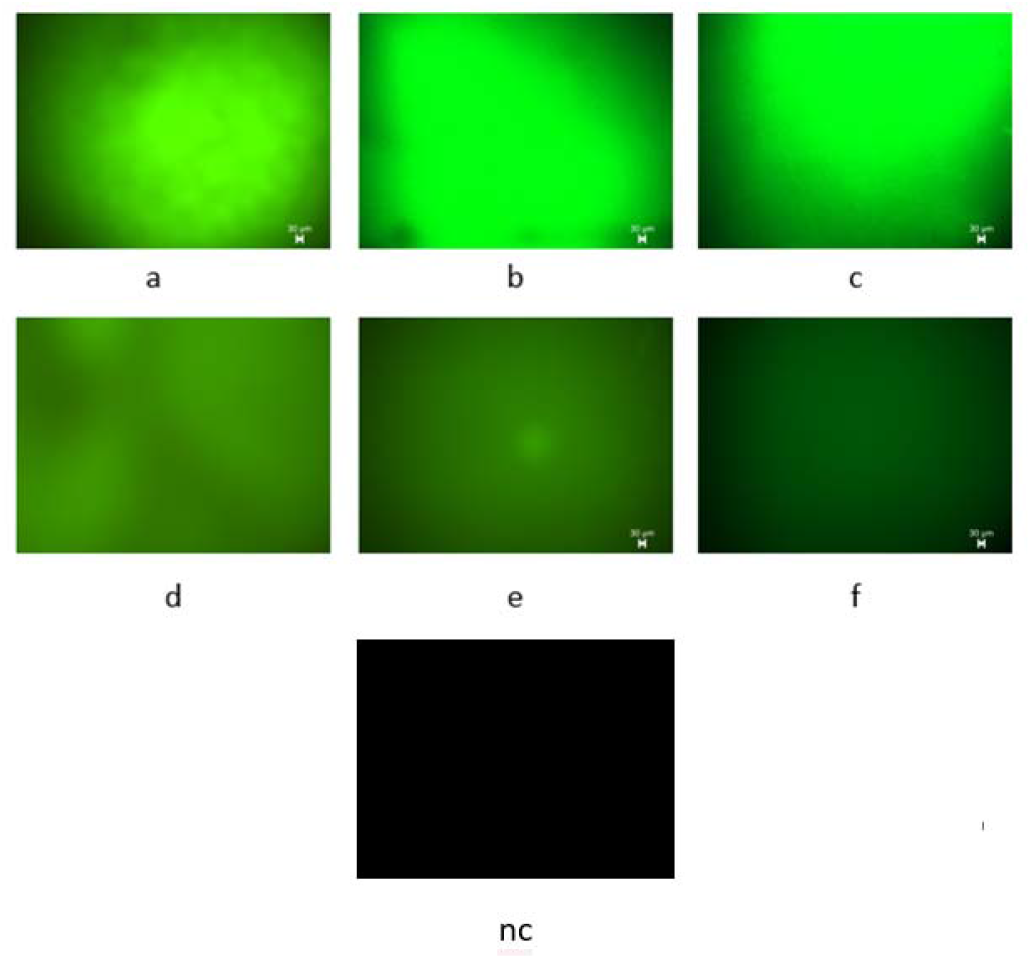
The panels shown here represent the microscopic images of the fresh samples as observed under the fluorescent microscope: a. Pureblood, b. 10x diluted blood, c. 100x diluted blood, d. 1000x diluted blood, e. 10000x diluted blood, f. 100000x diluted blood, and nc. negative control. All the dilutions were made in PBS.

### Reaction with diluted blood samples

Fresh blood samples from 10x to 1,00,000x dilution levels were subjected to the reaction. A positive observation was made in each sample type.

The fluorescent spectra for aged diluted samples were also detected. The maximum age of the diluted sample to give positive results was found to be 17 days-minimum being six (6), where 1lakh times diluted sample failed to provide any signal (Figure 2 and 3).

**Figure 2:**
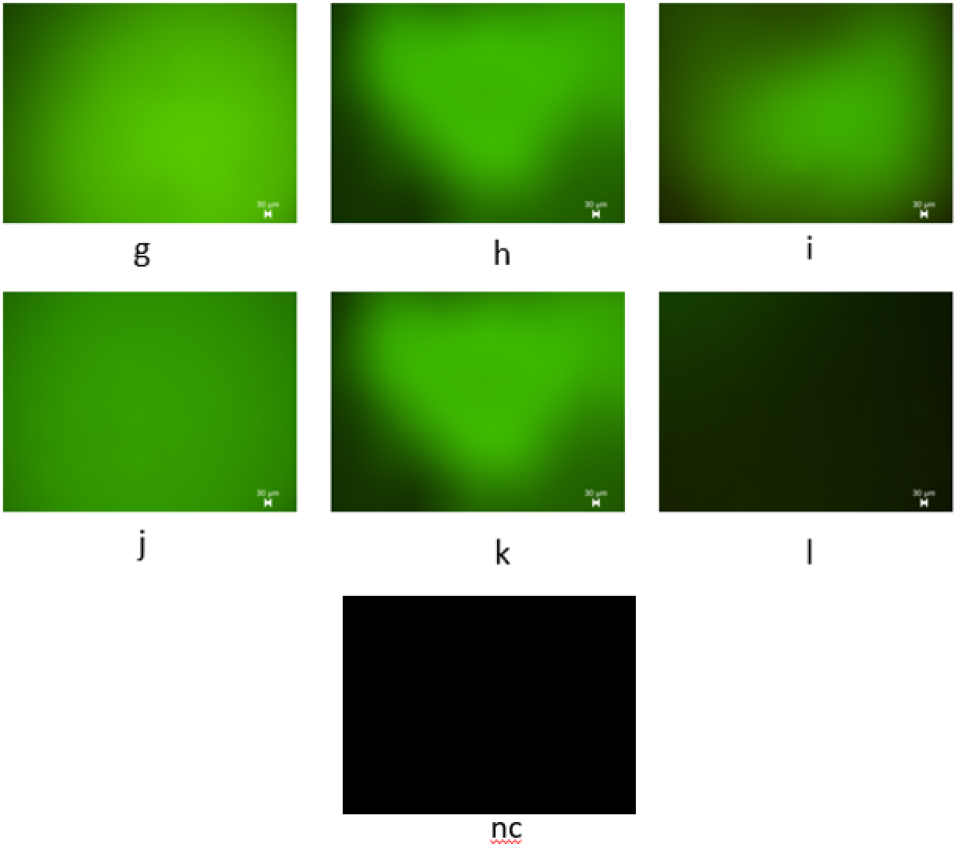
The panels represent the microscopic images of the 6-day old samples as observed under the fluorescent microscope: g. Pureblood, h. 10x diluted blood, i. 100x diluted blood, j. 1000x diluted blood, k. 10000x diluted blood, and l. 100000x diluted blood. Panel l shows unobtrusive fluorescence. nc as shown is a negative control.

**Figure 3:**
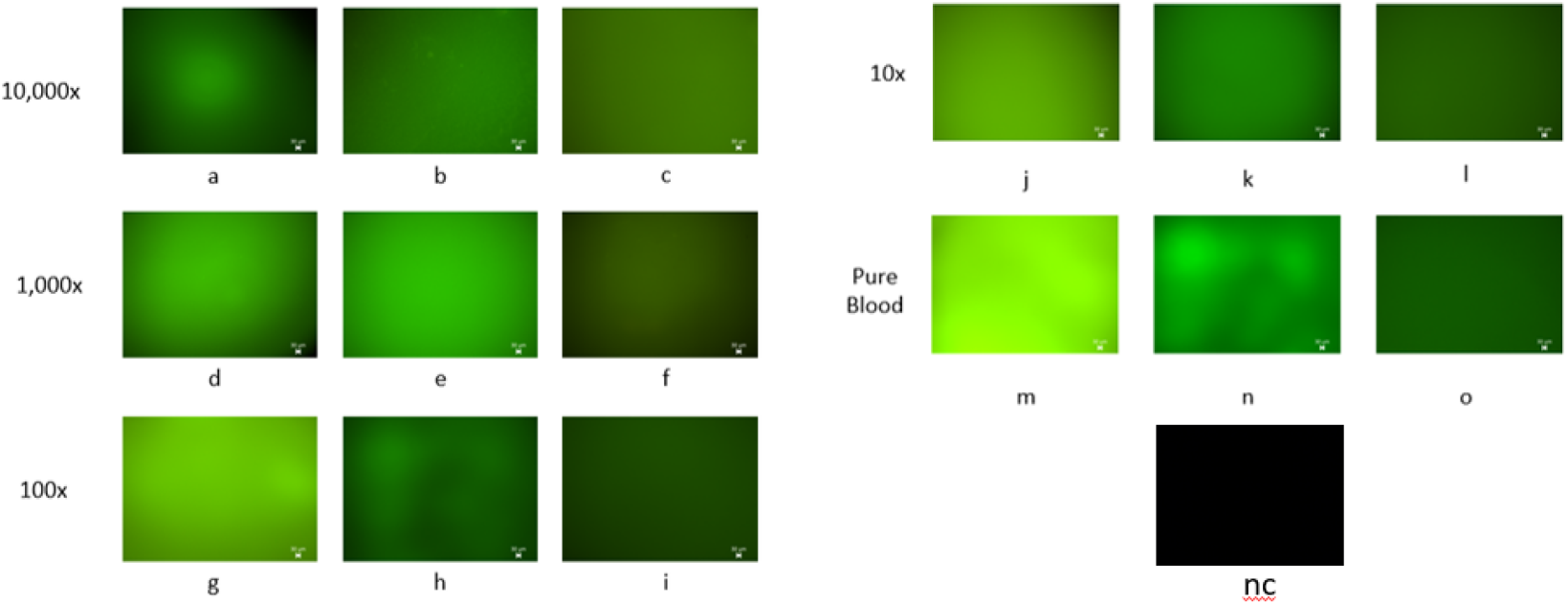
The panels a-o represent the comparative microscopic images of aged samples as observed under the fluorescent microscope. From a-c: 10,000x diluted sample aged 6,7, and 8 days; d-f: 1000x diluted samples aged 8, 10, and 13 days; g-i: 100x diluted samples aged 10, 15, and 17 days; j-l: 10x diluted samples aged 10, 15, and 17 days; and m-o: pureblood samples aged 10,15, and 20 days. 10,000x samples older than eight days didn’t show any fluorescence. Similarly, 1000x, 100x, and 10x lost their activity after 13, 17, and 17 days respectively. However, pureblood lost its fluorescent signal after 20 days of aging. nc represents the negative control.

Figure 4 summarizes the results, in days, pronounced from the samples. It is seen that the pure blood aged 20 days provides an apparent fluorescence, whereas the highly diluted samples can give positive results for up to a week.

**Figure 4:**
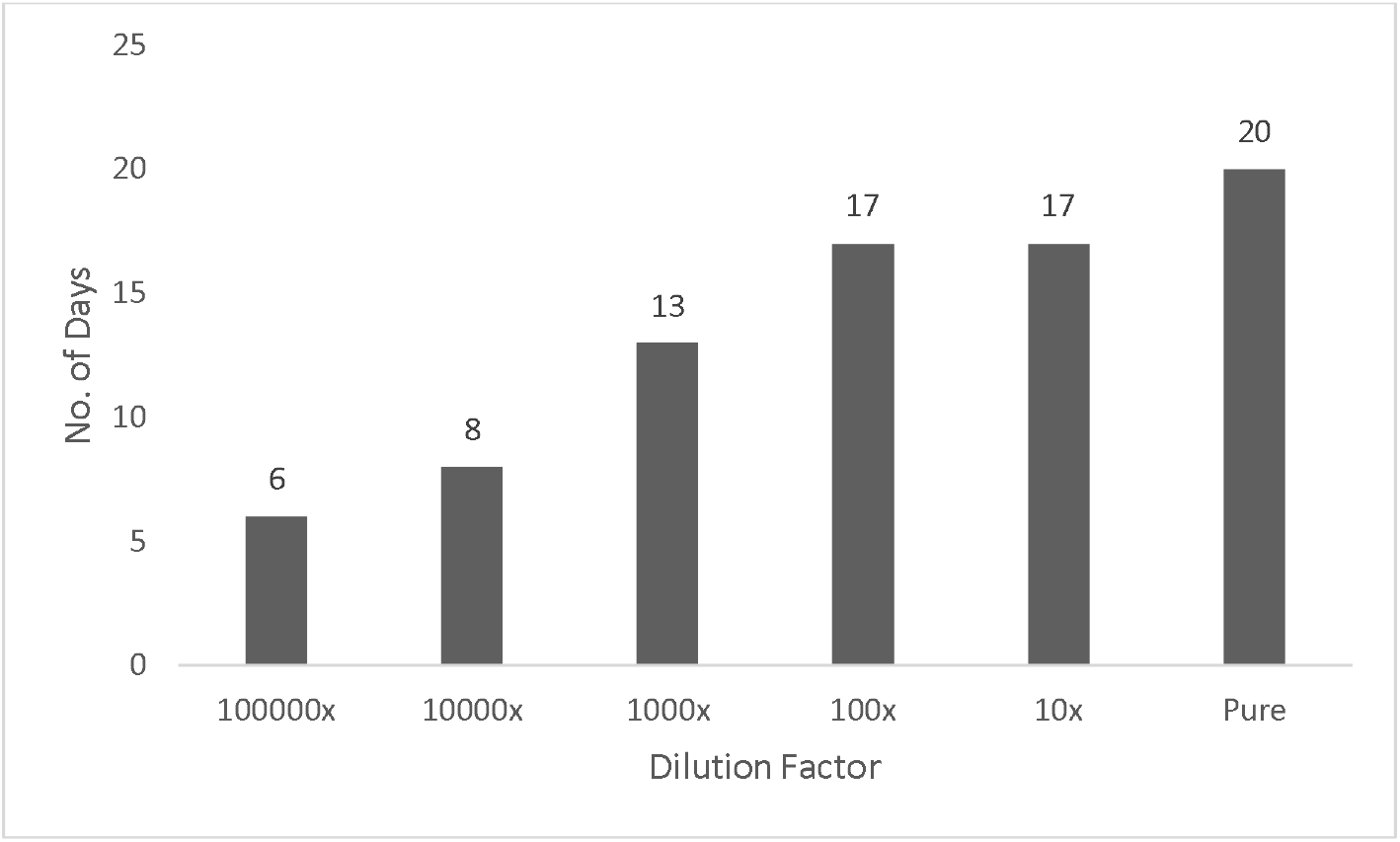
Graphical representation of the fluorescence observed in aged (number of days) diluted and pureblood samples.

### Reaction with Similar-looking stains

Beetroot juice, Tomato Ketchup, and Pomegranate juice were tested to see if they produced false-positive results against the proposed assay. It was reported that none of these materials reacted with the dye. Further, the stains were aged and tested again, but no fluorescent spectra were observed (figure 5).

**Figure 5:**
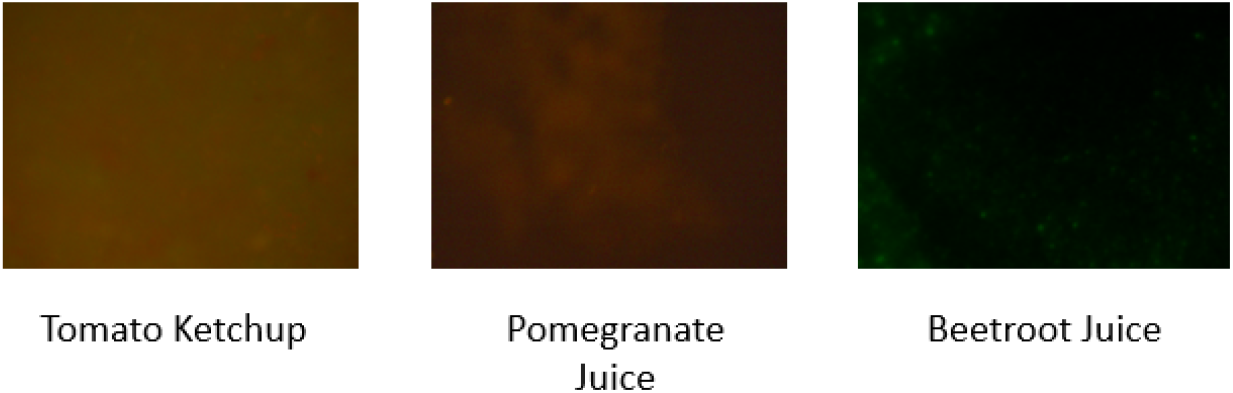
Fluorescent microscopic images of similar-looking stains showing no “glow.”

This defines that the proposed novel dye-based methodology has specificity toward blood and doesn’t tend to give false-positive results with other similar-looking materials.

## Discussion

In summary, the 2’,7’-dichlorofluorescein diacetate (DCFDA) dye has been systematically assessed for its efficacy in preliminary blood screening. DCFDA is a chemically reduced form of fluorescein. Researchers have been using it as an indicator for cells’ reactive oxygen species (ROS). The cell membrane permeable DCFDA is deacetylated inside the cells in the presence of a cellular esterase enzyme. It is then further oxidized to 2′,7′-dichlorofluorescein (DCF) via Fenton’s reaction if iron is present in the cellular milieu (Figure 6).

**Figure 6:**
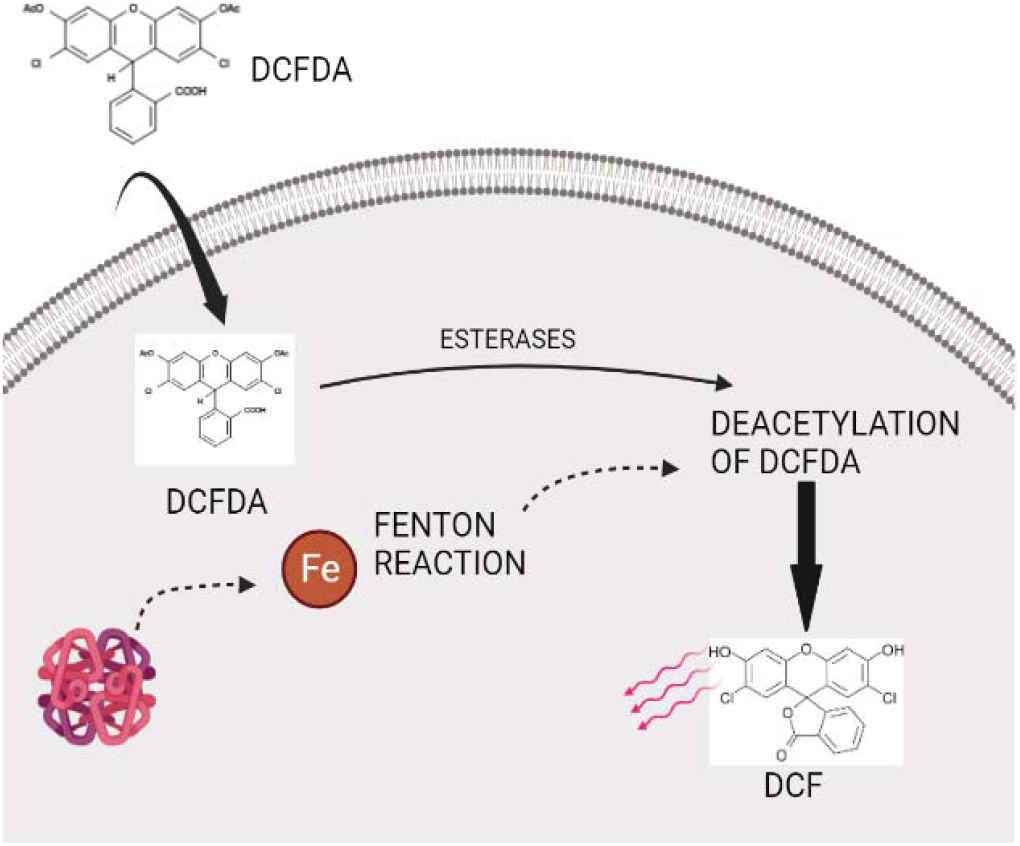
Transformation of 2’,7’-dichlorofluorescein di-acetate (DCFDA) inside the cell into Dichloro-fluorescein (DCF) via cellular esterases and iron moiety undergoing Fenton reaction. The DCFDA thus gets converted into DCF-a fluorescent dye.

Generally, hydrogen peroxide in the Fenton’s reaction oxidizes Fe^+2^ to Fe^+3^, producing hydroxyl radical and an anion. Again, the same hydrogen peroxide reduces Fe^+3^ to Fe^+2^, producing a peroxide radical and a proton. The reaction is favored at acidic pH. The reaction is as follows:

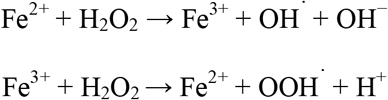

This DCF is fluorescent in nature and can be quantitatively detected either by fluorescence microscopy at excitation and emission wavelengths of 480 nm and 530 nm, respectively (or with excitation sources and filters suitable for fluorescein (FITC)) or using fluorometry.

The present study screened fresh and aged samples using the proposed DCFDA dye. It was further reported that the dye could positively detect 20-day-old bloodstains. Any preliminary test would be called the best fit when it is specific and sensitive. Thus, to test the assay’s sensitivity, diluted blood samples were subjected to the detection where the dye could screen 17 days old (100 parts diluted) blood samples. Additionally, it was also observed favorable results up to one lakh parts of dilution. Further to check the specificity of the assay, similar-looking stains, namely Pomegranate juice, Beetroot juice, and Tomato ketchup, were examined for their reaction with the said dye. None of these samples gave a positive fluorescence spectrum.

Hence in routine analysis, it would be beneficial to use DCFDA as an adequate replacement for all the frequently used blood screening reagents in forensic laboratories. It will avoid the possible chronic exposure to hazardous reagents like benzidine, phenolphthalein, etc. The false-positive results encountered using these reagents may hinder the investigation. Still, the results from the study show that DCFDA does not give false-positive results with the substances tested. However, a diverse group of materials may show false-positive results when reacted with DCFDA. Since this list could be endless, possible reactants were tested, and the results were satisfactory, demonstrating the specificity of the dye to blood.

## Conclusion

Primary blood screening assays are the method of choice by forensic experts to test the presence of blood at a crime scene. These assays are based on the redox reactions between the reagent and the blood hemoglobin in the presence of peroxide. There have been several studies reporting the ill effects of these chemicals. However, no reports have yet been reported on their replacement. In the present study, DCFDA-a non-carcinogenic, non-toxic dye, has been tested for its specificity and sensitivity against the blood specimen. It has been observed that the dye distinctly detects fresh and aged blood samples at minimal concentrations for up to 20 days. Conventionally used reagents have additional limitations, showing false positive reactions with samples like vegetable and fruit juices. To testify this, DCFDA has been tested against a few similar-looking samples like tomato ketchup, pomegranate and beetroot juice, where no reaction was observed. Thus, it is proposed that the fluorescein-derived DCFDA dye can be an excellent alternative to the currently used assays for blood identification.

